# Predictive eye movements anticipate target rebounds in an interception task

**DOI:** 10.1101/2025.03.17.643715

**Authors:** Mario Treviño, Nathaly Martín, Francisco Hernández, Andrea Barrera, Inmaculada Márquez

## Abstract

Predictive control in humans enables anticipatory behavior by combining sensory feedback with internal forward models. In interception tasks, such predictive processing could enable the visual system to estimate future target positions, thereby facilitating precise and timely motor responses. This study investigated the existence of predictive fixations during a visuomotor task where participants used a joystick to intercept a target that rebounded within a circular arena. We categorized eye movements into fixations, smooth pursuits, and saccades using a threshold-based classification method. The task’s circular geometry simplified trajectory analysis by consistently redirecting the target toward the arena’s center after rebounds, enabling clear identification of predictive fixations. Results showed that participants consistently aligned their gaze and joystick movements with the target’s anticipated trajectory before rebounds. Similarly, fixation and gaze onsets showed pre-rebound adjustments, using learned statistical regularities of rebounds to anticipate and prepare for changes in the target’s trajectory. Directional cues from the target’s entry angles just before rebounding influenced gaze alignment and prediction accuracy, while target speeds affected fixation locations. Moreover, masking the target disrupted gaze alignment and increased variability, whereas masking the user had minimal effects, highlighting the importance of target visibility in predictive control. These findings demonstrate the crucial role of predictive fixations in visuomotor coordination, offering a broader understanding of anticipatory behaviors and their applications in dynamic tasks. Our task’s design offers a robust framework for studying predictive processing across individuals, with implications for sports performance, clinical diagnostics, and human-computer interaction.

## INTRODUCTION

Eye movements and fixations are crucial for investigating how sensory predictions coordinate with motor responses, particularly in tasks based on achieving specific goals. Fixations (periods when the gaze is steadily held on a particular location) serve as "anchors" for visual attention, providing essential visual information. In contrast, saccades (rapid eye movements between fixations) enable quick gaze shifts with minimal sensory input. During saccades, humans experience "saccadic suppression," a phenomenon that reduces visual sensitivity and prevents motion perception, thereby maintaining visual stability. The brain implements a variety of mechanisms, such as efference copy signals, to maintain perceptual stability by anticipating saccades and adjusting spatial coordinates accordingly. These strategies ensure continuous perception across saccades by continuously integrating visual information before and after these rapid eye movements.

Anticipatory saccades can enhance performance by positioning the eyes in task-relevant areas before action completion, facilitating efficient feedback processing for error correction and motor adjustments (Brand et al. 2024). Rooted in the framework of forward models (where the brain is proposed to perform Bayesian inference requiring a generative model to predict observations), eye movements are believed to depend on the central nervous system’s internal representations that predict sensory consequences of motor actions. Such predictions enable error detection by comparing expected and actual outcomes (Joch et al. 2017). In line with these ideas, the ideomotor theory posits that actions are initiated based on anticipated sensory outcomes, serving as a basis for intentional, goal-oriented movements and agency (Koch et al. 2004; Pfeuffer et al. 2016).

The concept of active vision further extends these ideas, emphasizing the role of gaze control in prioritizing regions of high informational value based on task demands and predictions, rather than solely relying on explicit, ‘bottom-up’ visual properties (Itti and Koch 2000). This mechanism enables individuals to direct their fixations toward areas critical for task success, facilitating the intake of relevant information (Henderson 2017). Thus, the premise suggests that predictive saccades and fixations enable the visual system to utilize internal models, aligning the gaze with anticipated stimuli to enhance visuomotor coordination. From this viewpoint, gaze control reflects a dynamic interplay of rapid saccades and stable fixations, guided by prior experience, familiar environmental patterns, and task-specific knowledge. Predictive saccades direct gaze toward objects seconds before interaction, facilitating effective spatial and temporal planning (Mennie et al. 2007). These eye movements often occur just before expected action outcomes, reflecting a proactive approach by the visual system to predict sensory events and acquire feedback promptly (Pfeuffer et al. 2016). Such behaviors illustrate the integral role of cognitive processes in shaping predictive fixations by allowing the visual system to prioritize relevant areas of interest and anticipate the next sequence of events.

Predictive fixations are essential for sensorimotor coordination, as they illustrate how the brain combines sensory input with motor actions. Their significance becomes even more pronounced in dynamic settings, where stimuli are in motion, and rapid adjustments are necessary. In activities like sports, this interplay between sensory anticipation and motor execution is crucial for success. For example, athletes in sports such as cricket or table tennis use predictive saccades to track the fast-moving ball effectively. By preemptively aligning their gaze with trajectory-critical points—such as the ball’s release or bounce— they achieve optimal timing and accuracy in their responses (Land and McLeod 2000). Such anticipatory alignment is proposed to rely on internal forward models and ideomotor anticipations, which enable the real-time detection and correction of errors. Therefore, predictive mechanisms probably enhance performance across a wide range of goal-directed activities by integrating sensory predictions with motor actions to ensure precision and efficiency (Brand et al. 2024; Mann et al. 2019; Shalom et al. 2011).

Building on these ideas, we recently developed a visually guided task to examine how individuals chase a moving target. The task required participants to control a white dot via a joystick to intercept a black dot (the target) within a circular arena. The target moved along rectilinear trajectories with variability introduced by changes in speed and direction, creating dynamic and uncertain task conditions (Treviño et al. 2024). In these experiments, we recorded performance metrics, including collision times, alongside eye-tracking measures to capture gaze properties such as gaze speed, gaze-to-target distance (GTD), and pupil dilation. Our initial findings identified task parameters that influenced gaze patterns, pupillary responses, and overall task performance (Márquez and Treviño 2024a, 2024b). However, a critical but unexplored aspect of this task involves characterizing gaze behavior specifically during target rebounds. In our task design, the target bounced off the arena walls according to the law of reflection (*i.e.*, the reflection angle equaled the incidence angle). Additionally, the rebounding angle included variability through a specified angular range for each trial (Treviño et al. 2024). Consequently, the rebounds presented participants with a particular challenge: to either anticipate the post-rebound trajectory or react to the target’s directional change after it rebounded.

Given the integral role of the oculomotor system in producing predictive fixations, we hypothesized that participants solving our task would use predictive gaze strategies to deal with these frequent rebound events. More specifically, we posited that predictive saccades and fixations occur before rebounds, reflecting proactive adjustments in anticipation of the target’s post-rebound trajectory. The current study examines oculomotor responses associated with target rebounds to evaluate the presence and extent of a predictive component in gaze behaviors. Our task and methods could be particularly relevant for understanding visuomotor coordination in critical settings, such as competitive gaming or medical training, where rapid and precise fixations on key visual elements are crucial for success.

## MATERIALS AND METHODS

### Participants

Building on data from previous experiments (Márquez and Treviño 2024b, 2024a), this study involved 103 right-handed adults (53 women) aged 18–29 years (mean age: 21.56 ± 0.20 years, mode: 22). Experiments were performed during 2023 using a within-subject design to assess behavioral responses across different conditions, such as varying target speeds and visual occlusion scenarios. Participants were divided into three experimental groups designed to investigate different aspects of predictive fixations and gaze behavior during the visuomotor interception task. The experiments are presented separately in different sections and figures as follows. Group E_1_: Forty participants completed 900 trials with a target moving at a constant speed of *v_T_* = 30°/s. Directional uncertainty was manipulated by varying the angular range (AR) from 0° to 180° while maintaining a fixed direction interval (FDI) of 100 ms (Márquez and Treviño 2024a). This experiment examined the predictive nature of fixations before target rebounds, also focusing on peri-event time histograms (PETHs) of gaze onsets to determine whether systematic changes occurred before and after rebounds. Additionally, we assessed the influence of entry and exit angles at the rebound point on fixation distributions, analyzing their orientation and variability before and after the rebound. Group E_2_: Thirty-three participants completed 900 trials, with *v_T_* ranging from 10°/s to 60°/s in increments of 10°/s, while AR was fixed at ±75° and FDI at 500 ms (Márquez and Treviño 2024b). With this experiment, we investigated how *v_T_* influenced fixation distributions: whether the center of mass of these distributions shifted with speed and how speed modulated fixation distribution variability. Group E_3_: Twenty participants performed 800 trials incorporating visual occlusion of either the target or participant feedback under controlled conditions (*v_T_* = 30°/s, FDI = 500 ms, AR = ±75°; (Márquez and Treviño 2024b). This experiment examined the impact of masking either the target or the user at different inter-dot distances, evaluating how occlusion affected fixation distributions before and after rebounds. We used quota sampling to ensure balanced gender representation. To be eligible, participants were required to have normal or corrected-to-normal vision, confirmed by the Snellen test, and report no history of psychiatric, neurological, or neurodevelopmental disorders. Participation was voluntary, without financial incentives, and all procedures adhered to ethical guidelines approved by the Instituto de Neurociencias ethics committee (ET102021-330 and ET122023-382). Written informed consent was obtained from all participants, and data confidentiality was maintained in accordance with institutional and ethical guidelines.

### Collecting demographic information from participants

We gathered demographic information before conducting the visuomotor tasks to ensure balanced representation and adherence to inclusion criteria. Participants provided self-reported data on age, gender, and socioeconomic status through open-ended questionnaires, which included clear written instructions to ensure consistency in their responses. We maintained participant confidentiality in accordance with institutional ethical standards for handling sensitive information. This demographic data collection helped us verify the absence of some confounding factors, such as age or gender differences, which could impact visuomotor performance, thereby contributing to the study’s robustness and generalizability.

### Visuomotor task

We utilized a visuomotor interception task where participants maneuvered a joystick to intercept a moving target. We designed the task to deliberately alter target motion dynamics, such as speed and directional variability, to introduce controlled uncertainty (Márquez and Treviño 2024b; Treviño et al. 2024). Participants were seated approximately 70 cm from a 27-inch monitor (1920 × 1080 pixels, 60 Hz). The task occurred in a circular arena on a gray background projected on a monitor screen, featuring both a user-controlled white dot and a computer-controlled black dot, each represented by filled circles with a visual angle of 0.3° (**Figure 1A**). At the start of each trial, the target appeared at random locations along the arena’s perimeter and traveled at constant speeds for the duration of the trial, ranging from 10°/s to 60°/s. We implemented directional changes for the moving target at fixed direction intervals (FDI) of 500 ms, creating predictable windows for trajectory changes. Target motion variability was controlled by adjusting the target’s angular range (AR), which defined the extent of potential changes in angular directions (Treviño et al. 2024). When the target hit the circular arena limits, it rebounded according to the law of reflection, with its post-bounce path additionally influenced by the specified AR (center and right panels in **Figure 1A**). Trials ended either when the participant successfully intercepted the target or after a 5-second time limit. Auditory feedback indicated trial outcomes (Treviño et al. 2021). Participants had to return the joystick to the center position before beginning the next trial. Each session included up to 900 trials, with two 5-minute breaks to minimize fatigue (Márquez and Treviño 2024b; Treviño et al. 2024). In experiments involving masking conditions, we measured how participants’ predictive responses depended on visual information. During specific trials, we temporarily masked either the user’s or target’s dot by setting their contrast to 0%, rendering them invisible. This occlusion was applied at different randomly permuted inter-dot-distances (IDD). The masking distances corresponded to IDD values of 0°, 0.8°, 1.4°, and 2.7°, respectively (Márquez and Treviño 2024b). The task was implemented in MATLAB R2023a using the Psychophysics Toolbox (PTB-3).

**Figure 1.**
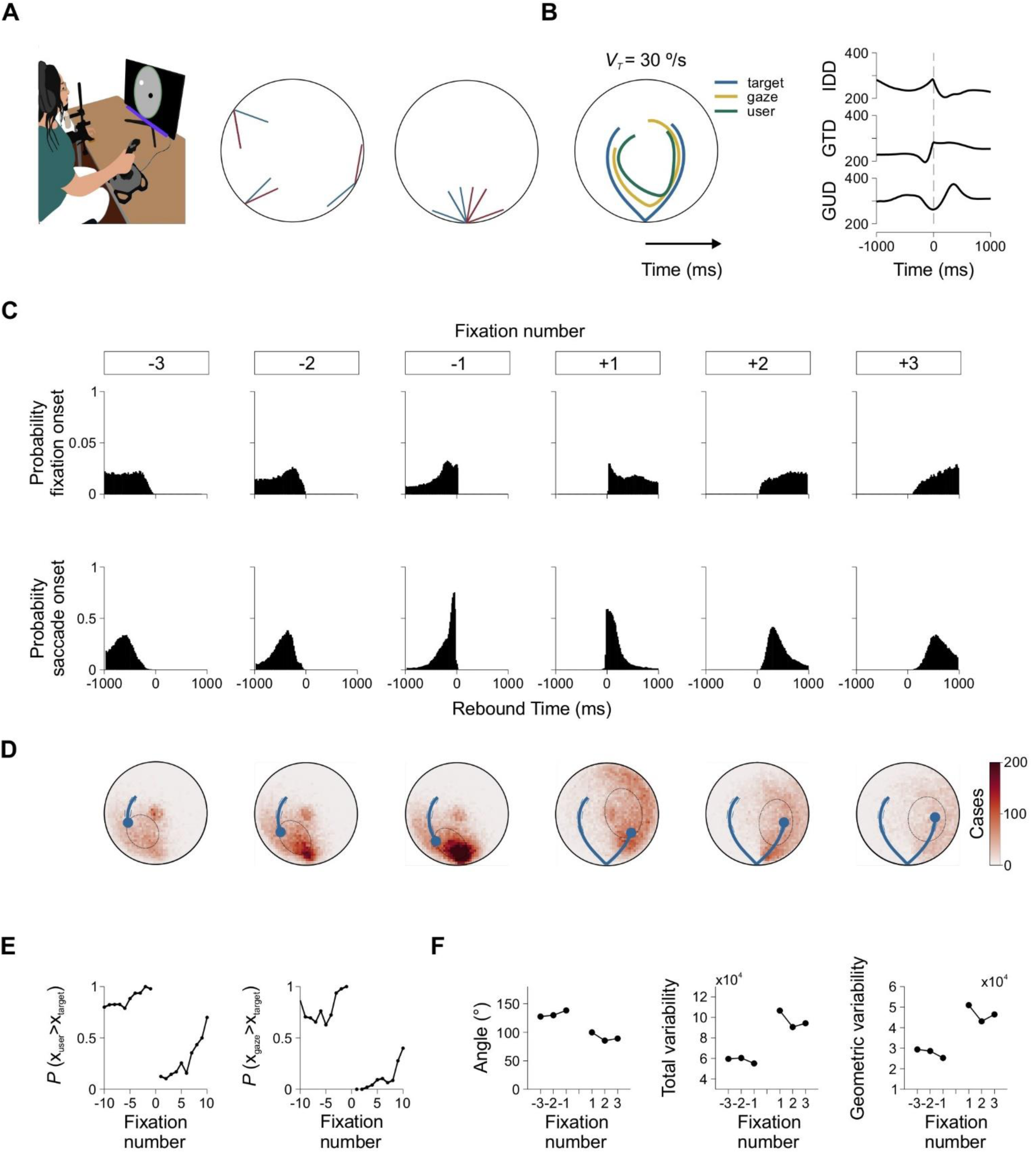
Predictive Fixations Around Target Rebounds. *A*: Participants used a joystick to control a white dot to intercept a black moving target within a circular arena. Insets illustrate potential rebound locations along the arena’s circumference. The right panel shows target rebounds aligned such that all rebounds enter from the left and exit to the right, (standardizing trajectory analysis). *B*: Mean trajectories of the target (sky blue), gaze (gold), and user (green teal), with portions to the right representing future times relative to the rebound (time zero). Right insets display event-triggered averages for inter-dot distance (IDD), gaze-to-target distance (GTD), and gaze-to-user distance (GUD), revealing distinct changes around rebounds. *C*: Peri-event Time Histograms (PETHs) for fixation onsets (upper row) and saccade onsets (lower row) show temporal alignment of eye movements relative to fixation numbers. Distributions shift closer to rebound times, reflecting gaze adjustments. *D*: Heatmaps depict fixation densities for discrete events before and after rebounds, with blue lines indicating the average target trajectory until each fixation. Fixation densities to the right of the target’s position suggest predictive behavior. *E*: Probabilities of joystick and gaze positions being to the right of the target increase steadily before rebounds and drop sharply afterward, reflecting predictive strategies. *F*: Average angles (left panel) and variability measures (total and geometric, center and right panels) exhibit robust changes around rebounds, highlighting shifts in spatial alignment and precision of gaze behavior.

### Gaze tracking

We recorded binocular eye movements using a Tobii eye tracker operating at 60 Hz, positioned below the monitor. Participants were stabilized with a chin rest and forehead support to minimize head movements, while ambient lighting was maintained at approximately 100 lux using curtains (Márquez and Treviño 2024a). The calibration process consisted of four steps: 1) a standard 4-point calibration using the eye tracker software, with fixation points located at the corners of the screen; 2) a custom MATLAB routine, where participants fixated on white dots (0.6° visual angle) positioned along the arena’s perimeter; 3) smooth pursuit measurement, where participants tracked a dot moving at 10°/s to assess gaze position and gaze gain (*G_G_* = *v_G_/v_T_*); 4) assessment of the pupillary light reflex through alternating black-and-white screens presented in 35-second trials (six cycles, repeated four times).

### Data analysis

We processed our data using MATLAB R2023a. Missing data, blink intervals, and outliers were interpolated linearly. We analyzed participants’ joystick movements by calculating the inter-dot distance (IDD), defined as the Euclidean distance between the user-controlled dot and the target, measured on the screen in either pixels or degrees of visual angle. User speed (*v_U_*, in °/s) measured the velocity of joystick movements on the screen, revealing how participants adapted to the target speed. The joystick gain was set to one, ensuring a linear relation between its tilt and the pixel movement of the white dot on the screen, and allowing the joystick’s position to reach the screen’s boundaries. We also extracted gaze positions and calculated gaze speed (*v_G_*, in °/s). Gaze-to-target distance (GTD) measured the Euclidean distance between the gaze position and the target on the screen, indicating visual tracking accuracy (Márquez and Treviño 2024b). To analyze responses to target bounces, we identified bounce events by detecting when the target’s radius touched the arena’s boundary radius. A ±60-frame window was employed to capture behavioral responses associated with each event, encompassing 1000 ms before and after the rebound. We then calculated entry and exit bounce angles along the arena’s circumference using a 3-frame interval (approximately 50 ms) to estimate directional vectors. We transformed all traces (gaze, joystick, and target trajectories) into polar coordinates relative to the arena center. All clipped traces were further standardized by aligning each bounce event to a common reference angle positioned at the bottom of the arena (**Figure 1B**). Only "complete" and "clean" bounces with high-quality trajectory data were included.

Using a dual-threshold method, we classified eye movements into saccades, smooth pursuits, and fixations (Liversedge et al. 2011). Saccades were identified as gaze movements with velocity (*v_G_*) ≥ 30°/s and acceleration (*a_G_*) ≥ 50°/s² (second derivative of position), with subsequent frames (100 ms) marked as part of the saccade. Smooth pursuit was detected in portions of gaze traces not classified as saccades, with *v_G_* ≥ 5°/s and velocity differences of ±20°/s relative to the target speed (*v_T_*), sustained for ≥ 50 ms. Finally. we defined fixations as gaze positions with a velocity *v_G_* < 3°/s that were not part of saccades or smooth pursuits.

To analyze the spatial distribution of fixations relative to the bouncing target, we generated heatmaps to describe fixation density. We extracted the mean vector (***m***) and covariance matrix (***C***) to quantify the central location and spread of fixation distributions. The mean vector ***m*** = [*m_x_*, *m_y_*] represents the center of the fixation distribution, while the covariance matrix ***C*** describes its spread across the horizontal (*x*) and vertical (*y*) dimensions:

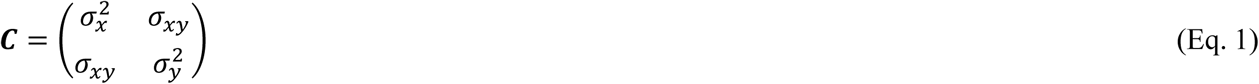

where *σ*_*x*_^2^ and *σ*_*y*_^2^ represent the variances along the *x* and *y* axes, and *σ_xy_* is the covariance. In some cases, we fitted and plotted 2D Gaussian ellipsoids around ***m*** at one standard deviation to capture the distribution’s shape and orientation. The orientation of the ellipsoid was determined by performing an eigendecomposition on ***C***:

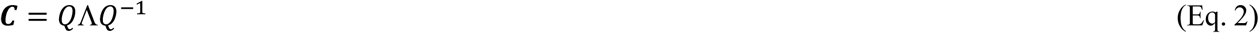

where *Q* contains the eigenvectors of ***C***, defining the principal axes of the ellipsoid, and Λ is a diagonal matrix with the corresponding eigenvalues, *λ_1_* and *λ_2_*. The eigenvector associated with the largest eigenvalue *λ_1_* indicates the direction of the ellipsoid’s major axis. Consequently, the angle *θ* of the major axis relative to the *x*-axis was calculated as:

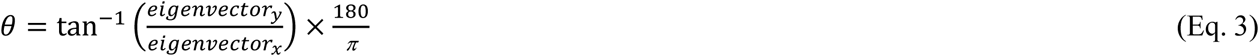

where *eigenvector_x_* and *eigenvector_y_* are the components of the primary eigenvector. Additionally, we derived two variability metrics from the covariance matrix ***C***. The total variability was calculated as the trace of ***C***, which sums the variances along each axis:

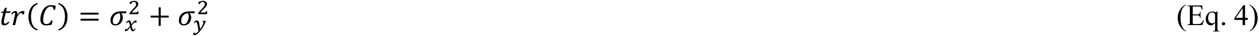

This metric reflects the overall dispersion of fixations in the *x* and *y* directions. Additionally, we calculated the geometric variability as the geometric mean of the eigenvalues of ***C***, offering an orientation-invariant measure of spread:

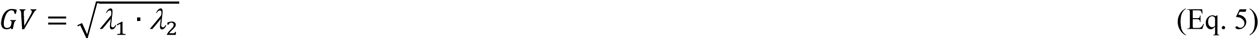

Throughout this manuscript, we operationally defined predictive behavior as the capacity to anticipate future target positions (Márquez et al. 2024). Positioning accuracy was assessed using squared distance errors (SDE, in px² or deg²), derived from the Euclidean distance between gaze and manual positions relative to current and future target positions. A 25-frame forward time window (∼416.66 ms) was utilized to evaluate predictive behaviors. The difference between future and current SDEs (ΔSDE) served as a metric for distinguishing more predictive (negative ΔSDEs) from more reactive (positive ΔSDEs) strategies (Brand et al. 2024; Brenner and Smeets 2011; Márquez and Treviño 2024b; Treviño et al. 2024).

### Statistical Analysis

To determine an appropriate sample size for our experiments (*i.e.,* sample size determination), we first estimated effect sizes from pilot data (not included in the current analyses). Cohen’s *d* values for paired comparisons of predictive fixation positions (paired t-tests) ranged from 0.91 to 5.27, indicating large to very large effects. Using G*Power 3 (Faul et al. 2007), we conducted an *a priori* power analysis, which indicated that a minimum of 15 participants was required to achieve 95% statistical power for detecting these effects at *α* = 0.05. For within-participant comparisons, we employed paired t-tests and used Pearson’s correlation coefficient to evaluate relationships between continuous variables. For onset timing analyses, we employed permutation tests (1,000 iterations) to generate null distributions, against which we compared observed fixation counts. We used repeated measures ANOVA to compare participant means across conditions, reporting the *F*-statistic, degrees of freedom, *P*-values, and omega squared (*ω*^2^). Omega squared is an effect size measure in ANOVA, estimating the variance proportion in the dependent variable due to the independent variable(s). Values are interpreted as 0.01 (small), 0.06 (medium), and 0.14 (large). It uses the sum of squares from the ANOVA table and adjusts for sample size. We implemented Bonferroni corrections to control for Type I errors in *post-hoc* analyses. We report all results as mean ± standard error of the mean (S.E.M.), with statistical significance set at *α* = 0.05. Figure panels indicate the number of participants for each task.

## RESULTS

This section presents the results from a detailed analysis of gaze behaviors while solving our visuomotor interception task. We examined participants’ eye movements—specifically saccades, smooth pursuits, and fixations—to determine whether and how predictive strategies could explain target trajectories before and after rebound events. We focused on gaze classification, the spatial distribution of fixations, and the effects of task parameters, such as trajectory angles, target speed, and masking, on anticipatory gaze patterns.

### Fixation distributions before and after rebounds

In our task, participants maneuvered a joystick to control a white dot, aiming to intercept a black dot (the target) moving within a circular arena (**Figure 1A**). The target followed two-dimensional rectilinear trajectories with constant speed (*v_T_*), fixed angular range (AR, see **Methods**), and fixed-direction intervals (FDI), which introduced stable levels of directional unpredictability (Treviño et al. 2024). Upon reaching the arena’s edges, the target bounced off according to the laws of reflection (center and right panels in **Figure 1A**). The enclosed circular geometry of the task arena involved multiple rebounds per trial, allowing us to study how participants adjusted their gaze and manual responses in relation to the target’s trajectory (approximately 1306 ± 162 rebounds per participant per session, *n* = 10). With this information, we investigated whether participants proactively aligned their saccades and fixations in anticipation of the rebound (Anderson et al. 2008) or responded reactively to changes in trajectory after the rebound (Shelhamer and Joiner 2003).

To analyze rebound-related behaviors, we first examined gaze and manual data from 40 participants who performed this task (Márquez and Treviño 2024b). We aligned trajectories of the target, joystick, and gaze around the rebounds to a common reference system by rotating the coordinates of traces for each rebound, ensuring all rebounds entered from the left and exited to the right (see **Methods**; center and right panels in **Figure 1A**). This procedure standardized our analysis across all rebound events, allowing consistent comparisons. Furthermore, this setup ensured that positions to the right of the target represent future events, while those to the left denote past events, as the target moved at a constant speed within this reference system **(Figure 1B)**.

Mean trajectories for the target (sky blue), gaze (gold), and joystick (green teal) provide a visual summary of how closely participants’ movements aligned with the target’s path (**Figure 1B**). We evaluated spatial accuracy, visual tracking precision, and gaze-manual coordination around rebounds by analyzing inter-dot distance (IDD), gaze-to-target distance (GTD), and gaze-to-user distance (GUD; (Márquez and Treviño 2024b). The average traces for these parameters revealed distinct shifts associated with rebounds (right panels in **Figure 1B**). Interestingly, the increase in IDD and GTD at the moment of rebound suggests that participants may be waiting for the particle after the rebound, as these values increase temporarily, indicating that the particle moves farther from both the user and gaze positions.

Next, we examined fixation locations relative to the rebounds by labeling them numerically for each participant as occurring before or after the bounce. Specifically, we analyzed the temporal and spatial distributions of three fixations preceding (-3, -2, -1) and following (+1, +2, +3) each rebound. The peri-event time histograms (PETHs) in **Figure 1C** display temporal distributions of fixation and gaze onsets, respectively. These histograms reveal the progressive temporal alignment of gaze onsets with the rebound, as the means and peaks of these distributions shifted closer to the rebound, in sequential order.

Heatmaps of fixation locations overlaid on the target trajectory illustrate areas of low density (whitish), to high density (dark red), providing a spatial representation of where participants fixated relative to the target. Fixations were numbered sequentially, and blue traces represented the average target position until the corresponding fixation (**Figure 1D**).

These heatmaps show that participants directed their fixations ahead of the target’s path before rebounds, as indicated by the heatmap peak positioned to the right of the current target location for fixations occurring before the rebound (*e.g.*, pre-1; **Figure 1D**). Statistical analyses confirmed these observations. Repeated-measures ANOVA revealed that joystick (main effect of fixation number: *F*_5,282_ = 100.8, *P* < 0.001, *ω*^2^ = 0.64; *post hoc* Tukey’s tests: *P* < 0.001), and gaze (*F*_5,282_ = 302.4, *P* < 0.001, *ω*^2^ = 0.81; *post hoc* Tukey’s tests: *P* < 0.001) positions consistently aligned ahead of the target across fixation numbers.

These results indicate that both fixations and joystick positions were consistently aligned to the right of the target’s location before rebounds, highlighting strong anticipatory behavior. However, a significant shift in alignment probabilities was observed, with both fixations and joystick positions showing a marked decrease in predictive characteristics after the rebound (**Figure 1E**). To further analyze fixation distributions, we fitted ellipsoids to fixation distributions and extracted key parameters, including orientation angle, total variability, and geometric variability (see **Methods**). The analysis confirmed pronounced changes in these parameters around the rebound. For example, variability decreased before the rebound, indicating tighter spatial alignment, but increased sharply afterward, reflecting a return to reactive adjustments (**Figure 1F**). These findings demonstrate the participants’ application of predictive visuomotor strategies, effectively using spatial and temporal cues to anticipate the target’s rebounding trajectory.

### Contrasting predictive and reactive behaviors using squared distance errors to future target positions

To investigate the predictive nature of fixations around rebounds, we used squared distance errors (SDEs). SDEs provided an unsigned measure of the spatial error between participants’ fixation locations and the target’s positions, enabling us to assess alignment accuracy for both current and future target locations. Utilizing a 25-frame forward time window (∼416.66 ms), we calculated SDEs for these positions. We calculated their difference (ΔSDE = SDE_future_ - SDE_current_) to facilitate the detection of predictive behaviors. Negative ΔSDE values indicated predictive behaviors, where participants’ gaze or manual positions were more closely aligned with future target positions. Conversely, positive ΔSDE values reflected more reactive behaviors, characterized by larger distances to the target’s current and future locations (Márquez et al. 2024). As illustrated in **Figure 2A**, the ΔSDE for gaze-to-target distance (GTD) across a forward time window of 25 frames (horizontal axis) shows strong negative values indicative of a predictive strategy, with a local minimum occurring around 225 ms (forward frame 14). Statistical analysis revealed significant effects across all fixation numbers (repeated measures ANOVA test, main effect of fix nr, *F*_11,564_ = 226.8, *P* < 0.001, *ω*^2^ = 0.68; Tukey’s *post hoc* tests in FF_14_ against current traces; *P* < 0.001). Box plots in **Figure 2B** illustrate the group differences between current SDEs, represented by gray whiskers on the left, and future SDEs calculated 14 frames ahead of the current target position. This difference was corroborated using a two-sample Kolmogorov-Smirnov test (test statistic = 0.8576; *P* < 0.001; **Figure 2C**). These results suggest that participants adjusted their fixations both before and after the target rebound, achieving optimal squared distance errors (SDE) by anticipating the target’s position approximately 225 ms (14 frames) in advance.

**Figure 2.**
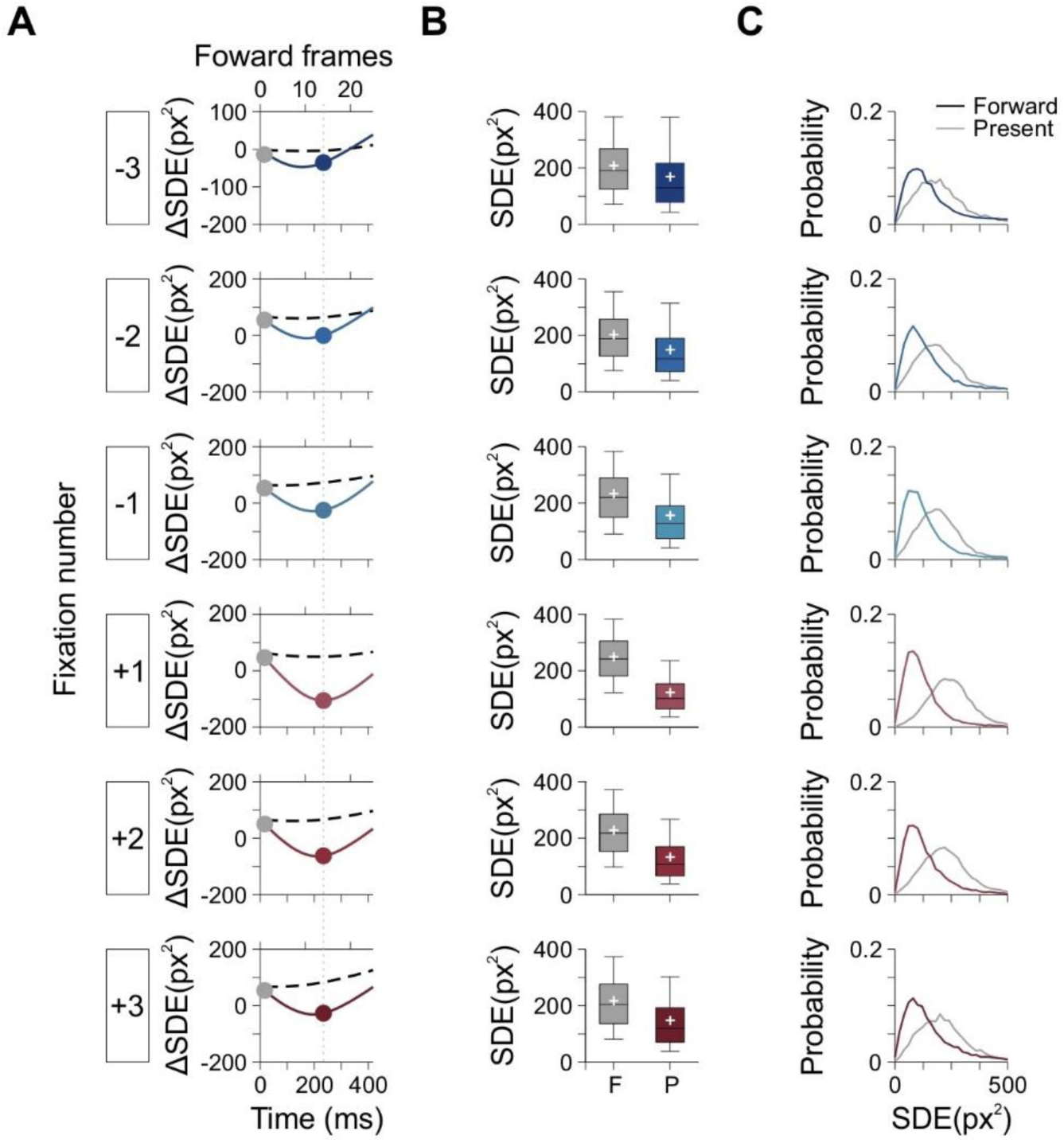
Predictive Behaviors Assessed via Squared Distance Errors (SDE). *A*: ΔSDE traces for fixations before and after rebounds across 25 forward frames. The x-axis shows forward frames (top) and corresponding temporal alignment (bottom). Negative ΔSDE values (downward trend) indicate predictive behavior, where responses align more closely with the anticipated future target position. Positive ΔSDE values (upward trend) would reflect reactive adjustments based on the target’s current position. A local minimum at forward frame 14 highlights the strongest predictive alignment. Black dotted lines represent ΔSDE values calculated using horizontally flipped gaze responses as a control. *B*: Whisker plots compare SDE values for current and future target positions. Gray whiskers represent observed SDE values for the current target position, while colored whiskers show future SDE values (forward frame 14) for each fixation. Reductions in SDE values across fixations reflect predictive adjustments. *C*: Probability distributions of SDE values for current (gray) and future (forward frame 14, colored) target positions. These distributions illustrate a systematic shift toward predictive strategies, with future SDEs showing greater alignment than current SDEs.

### Fixation and gaze onsets reflect predictive preparation before the target bounces

In sports involving a bouncing ball, fixation, and gaze onsets indicate predictive preparation before the target rebounds. These eye movements often occur before rebound points, demonstrating proactive visual attention and reliance on learned statistical regularities. Saccades similarly align with rebounds, driven by anticipatory neural activity in the occipitoparietal cortex. The premise is that predictive behaviors enhance performance and reduce reaction times by combining internal models and sensory cues (Badler and Heinen 2006; Gould et al. 2011). Strategic adaptations, such as look-ahead fixations and reduced blinking during critical moments, further highlight the proactive nature of predictive gaze control (Mennie et al. 2007; Shalom et al. 2011). Differences in saccade latency between successful and missed interceptions also suggest that predictive valuation processes guide saccades toward anticipated target locations, optimizing interception success (Brand et al. 2024; Mann et al. 2019).

Building on these ideas, we hypothesized that predictive processing during rebounds in our task would manifest as anticipatory shifts in fixation, and gaze onsets, aligning with expected rebound events and trajectory changes of the moving target. Specifically, fixation onsets were expected to adjust systematically before and after the rebound, reflecting proactive visual stabilization in preparation for the target’s rebound. Therefore, if predictive processing was indeed present, these fixation onsets should display systematic variations relative to the moment of the rebound. We collected the onset times of fixations, smooth pursuits, and saccades relative to the rebound event. We then constructed peri-event time histograms (PETHs) to measure occurrences within a time window from -600 to +600 ms around the rebound, divided into 10 ms bins. These event counts were then normalized into probabilities by dividing the frequency of cases in each bin by the total number of events within the analyzed window. To evaluate significance, we compared the observed probabilities of each eye movement type against probability distributions generated by permuting the event counts 1000 times. Probabilities were classified as significant (indicated in black) or not significant (shown in white) depending on whether the observed value fell below the 5th percentile of the permuted distribution tails (**Figure 3A**).

**Figure 3.**
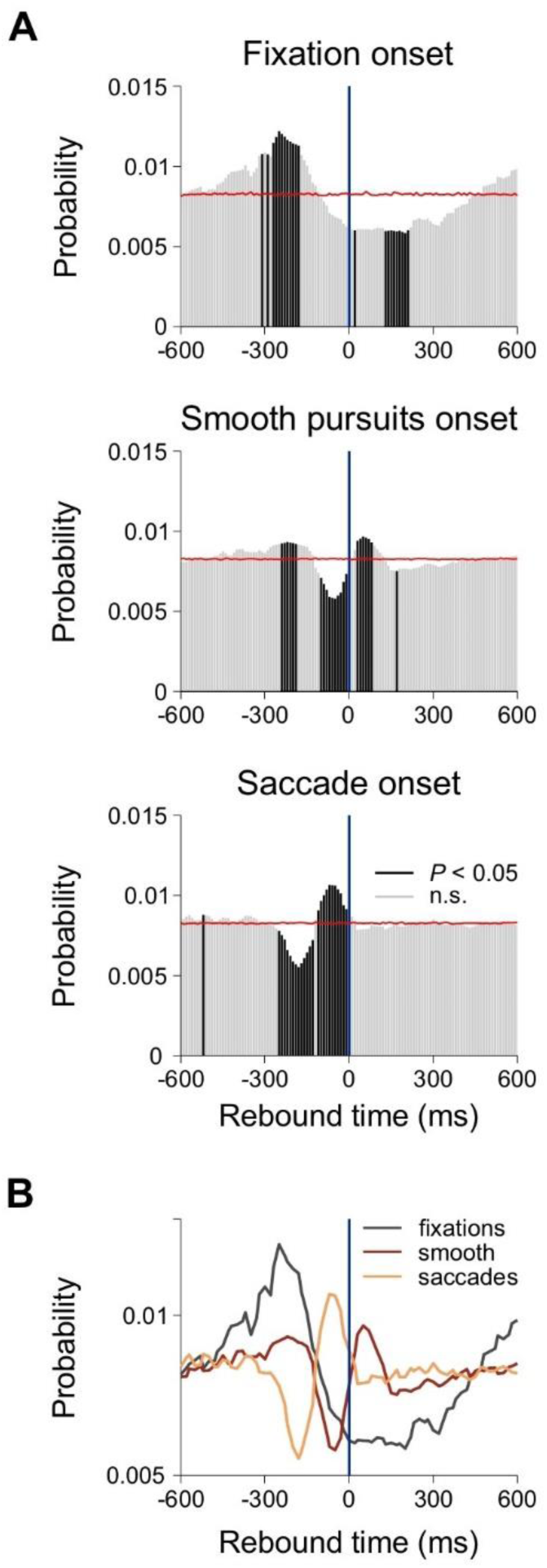
Temporal Dynamics of Fixation, Smooth Pursuit, and Saccade Onsets Around Target Bounces. *A*: Perievent time histograms (PETHs) display onset probabilities for fixations (top panel), smooth pursuits (middle panel), and saccades (bottom panel) within a ±600 ms window centered on target bounces. Bar plots show onset probabilities: significant values (black) are determined by comparing observed probabilities to a null distribution generated via random permutation of event sequences (1000 iterations), while non-significant values are shown in gray. Red traces indicate the mean probabilities derived from the permuted data, serving as the chance baseline. *B*: Overlay of onset traces for fixations (dark gray), smooth pursuits (dark red), and saccades (sandy brown) illustrates the temporal progression of gaze adjustments relative to the target bounce (time zero).

The PETHs reveal a distinct sequence of events, illustrated by the overlaid color-coded distributions shown in **Figure 3B**. Target rebounds were associated with an increase in fixation onsets before the rebound, followed by a slow decrease afterward (dark gray). In contrast, smooth pursuit (dark red) and saccade (sandy brown) onsets exhibited faster and opposing dynamics. Smooth pursuits showed a reduction in onset frequency before the rebound but increased afterward. Conversely, saccade onsets initially decreased but then surged just before the rebound, reflecting precise anticipatory adjustments. This temporal pattern highlights the interplay between anticipatory and reactive visuomotor behaviors around the rebound event.

### Influence of entry and exit angles on predictive fixations

The trajectory angles at which the particle enters the rebound point could provide directional cues shaping gaze alignment and influencing participants’ ability to predict the target’s future path. Entry angles define the particle’s trajectory as it approaches the rebound point, while exit angles describe its immediate direction after rebounding. Consistent and repeatable trajectories at these angles could facilitate the development of robust internal models, allowing participants to anticipate and adapt to task dynamics with greater accuracy (Bond and Taylor 2015; Coen-Cagli et al. 2009; Henriques et al. 1998; Sailer et al. 2005; Taylor and Ivry 2011). Conversely, unpredictable paths increase cognitive demands for recalibrating predictions, often resulting in higher error rates. Evidence from sports and motor learning supports these ideas. For example, expert volleyball players and skilled cursor-tracking participants utilize consistent trajectory cues to optimize their anticipatory responses (Piras et al. 2014; Sailer et al. 2005).

We categorized the entry and exit angles of the rebounding target into distinct ranges to investigate how these paths influence fixation behavior in our task. The entry angles were grouped into ranges of 170°±10°, 150°±10°, 130°±10°, 110°±10°, and 90°±10°, while the exit angles were categorized into 90°±10°, 70°±10°, 50°±10°, 30°±10°, and 10°±10°. Heatmaps of fixation distributions revealed spatial patterns that reflected participants’ alignment with the particle’s entry (sample panels on the left) and exit (sample panels on the right) trajectories (**Figure 4A**). Statistical analysis using repeated-measures ANOVA confirmed significant effects of fixation number on alignment across various entry angles (repeated measures ANOVA test, main impact of fix nr in forward trace #14 against current traces, entry angle = 170°: *F*_11,552_ = 103.5, *P* < 0.001, *ω*^2^ = 0.56; Tukey’s post hoc test, *P* < 0.001; entry angle = 150°: *F*_11,552_ = 145.8, *P* < 0.001, *ω*^2^ = 0.61; Tukey’s post hoc test, *P* < 0.001; entry angle = 130°: *F*_11,564_ = 177.4, *P* < 0.001; Tukey’s post hoc test, *P* < 0.001; entry angle = 110°: *F*_11,552_ = 130.4, *P* < 0.001, *ω*^2^ = 0.64; Tukey’s post hoc test, *P* < 0.001; entry angle = 90°: *F*_11,516_ = 102.4, *P* < 0.001, *ω*^2^ = 0.62; Tukey’s post hoc test, *P* < 0.001). Likewise, fixations consistently aligned with the target’s future trajectory across all exit paths, reinforcing their predictive nature (repeated measures ANOVA test, main effect of fix nr in forward trace #14 against current traces, exit angle = 90°: *F*_11,456_ = 59.14, *P* < 0.001, *ω*^2^ = 0.52; Tukey’s post hoc test, *P* < 0.001; exit angle = 70°: *F*_11,552_ = 140, *P* < 0.001, *ω*^2^ = 0.61; Tukey’s post hoc test, *P* < 0.001; exit angle = 50°: *F*_11,528_ = 154.4, *P* < 0.001, *ω*^2^ = 0.62; Tukey’s post hoc test, *P* < 0.001; exit angle = 30°: *F*_11,552_ = 170.7, *P* < 0.001, *ω*^2^ = 0.64; Tukey’s post hoc test, *P* < 0.001; exit angle = 10°: *F*_11,552_ = 126.3, *P* < 0.001, *ω*^2^ = 0.60; Tukey’s post hoc test, *P* < 0.001). Kolmogorov-Smirnov tests further validated these findings, showing significant differences in fixation distributions across all tested angles (*P* < 0.001 for all cases).

**Figure 4.**
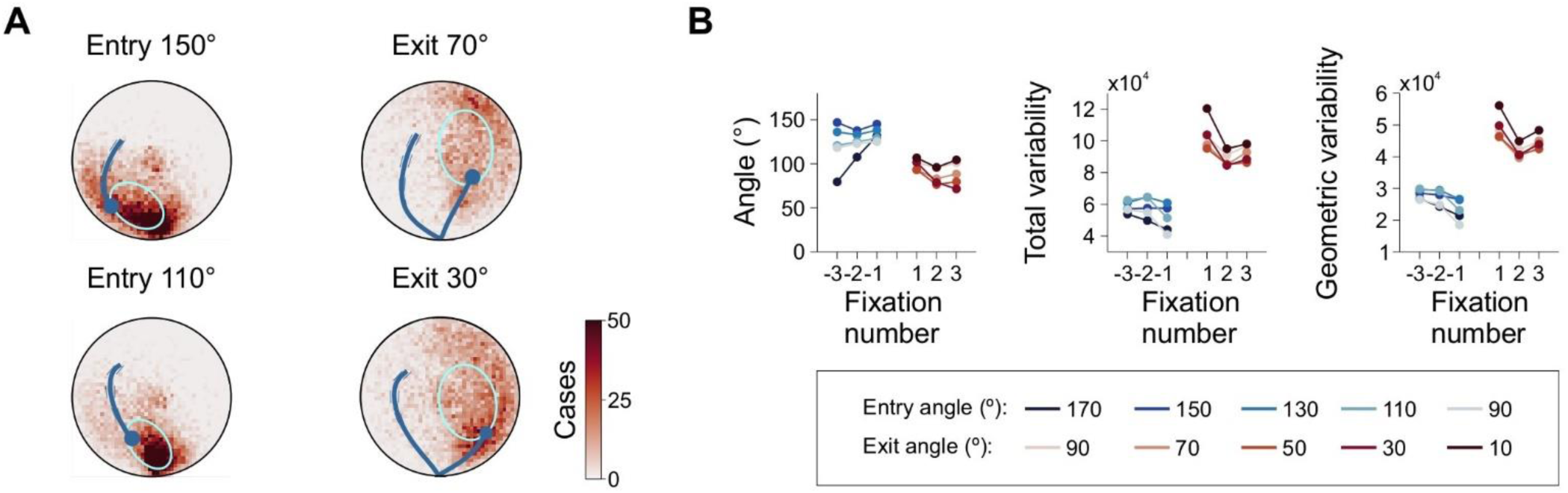
Influence of Entry Angles on Fixation Distributions After Rebounds. *A*: Heatmaps of fixation distributions for two example entry angles (left) and two example exit angles (right) overlaid on the target trajectory. The heatmaps represent fixation density, with colors ranging from low density (whitish) to high density (dark red). Ellipsoids fitted to the fixation distributions illustrate the spatial spread and orientation of fixations. *B*: Panel on the left shows average orientation of ellipsoids fitted for entry angles (blue colorscale) and exit angles (red colorscale) as a function of fixation number relative to the rebound. Center and right panels depict total variability and geometric variability of fixation distributions, respectively, across fixations.

We also examined how entry and exit angles affected the orientation of fixation distributions, which we characterized by measuring the angles of the major axis of Gaussian ellipsoids fitted to these distributions. Correlation analyses revealed no significant effect of entry angles on the distribution angles before rebounds (Pearson correlation, *r* = 0.579, *P* = 0.306; Spearman’s rank-order correlation, *ρ* = 0.700, *P* = 0.233). However, the initial fixations after rebounds did correlate with entry angles (*r* = -0.903, *P* = 0.018; *ρ* = -0.90, *P* = 0.042). In contrast, exit angles showed no correlation with fixation distributions before or after rebounds (*r* = -0.567, *P* = 0.319; *ρ* = -0.300, *P* = 0.683; left panel in **Figure 4B**).

Variance metrics of fixation distributions, including total and geometric variability (see **Methods**), confirmed the observation that entry angles, but not exit angles, influenced gaze behavior before rebounds (center and right panels in **Figure 4C**). Angles of post-rebound fixation distributions showed strong alignment with entry angles, indicating that directional cues before rebounds shaped predictive gaze behavior. In contrast, the lack of influence from exit angles suggests that post-rebound gaze adjustments relied primarily on pre-rebound information rather than exit direction. Therefore, target trajectory entry angles before rebounds influence anticipatory visuomotor responses to the rebounding target.

### Influence of target speed on predictive fixations

Target speed (*v_T_*) directly influences the time available for prediction and reaction, affecting participants’ ability to anticipate and respond to trajectory changes. Consequently, as *v_T_* increases, the cognitive and visual demands of the task also rise, potentially leading to shorter fixation durations and necessitating quicker saccades (Brand et al. 2024; Mann et al. 2019; Márquez and Treviño 2024b; Shalom et al. 2011; Treviño et al. 2024). We examined how *v_T_* influenced predictive responses to the bouncing target in our task. Our analysis revealed that the centers of mass of fixation distributions were impacted by changes in *v_T_* (**Figure 5A**). Repeated-measures ANOVA showed significant differences in ΔSDE (difference between future and current squared distance errors) across fixation numbers for speeds ranging from 10°/s to 60°/s (*v_T_* = 10 °/s, *F*_11,456_ = 29.17, *P* < 0.001, *ω*^2^ = 0.32; *v_T_* = 20 °/s, *F*_11,456_ = 72.44, *P* < 0.001, *ω*^2^ = 0.40; *v_T_* = 30 °/s, *F*_11,456_ = 87.32, *P* < 0.001, *ω*^2^ = 0.31; *v_T_* = 40 °/s, *F*_11,456_ = 89.78, *P* < 0.001, *ω*^2^ = 0.26; *v_T_* = 50 °/s, *F*_11,456_ = 74.41, *P* < 0.001, *ω*^2^ = 0.17; *v_T_* = 60 °/s, *F*_11,456_ = 54.81, *P* < 0.001, *ω*^2^ = 0.07). Consequently, participants positioned their pre-rebound and post-rebound fixations closer to the future positions of the target for all *v_T_* tested.

**Figure 5.**
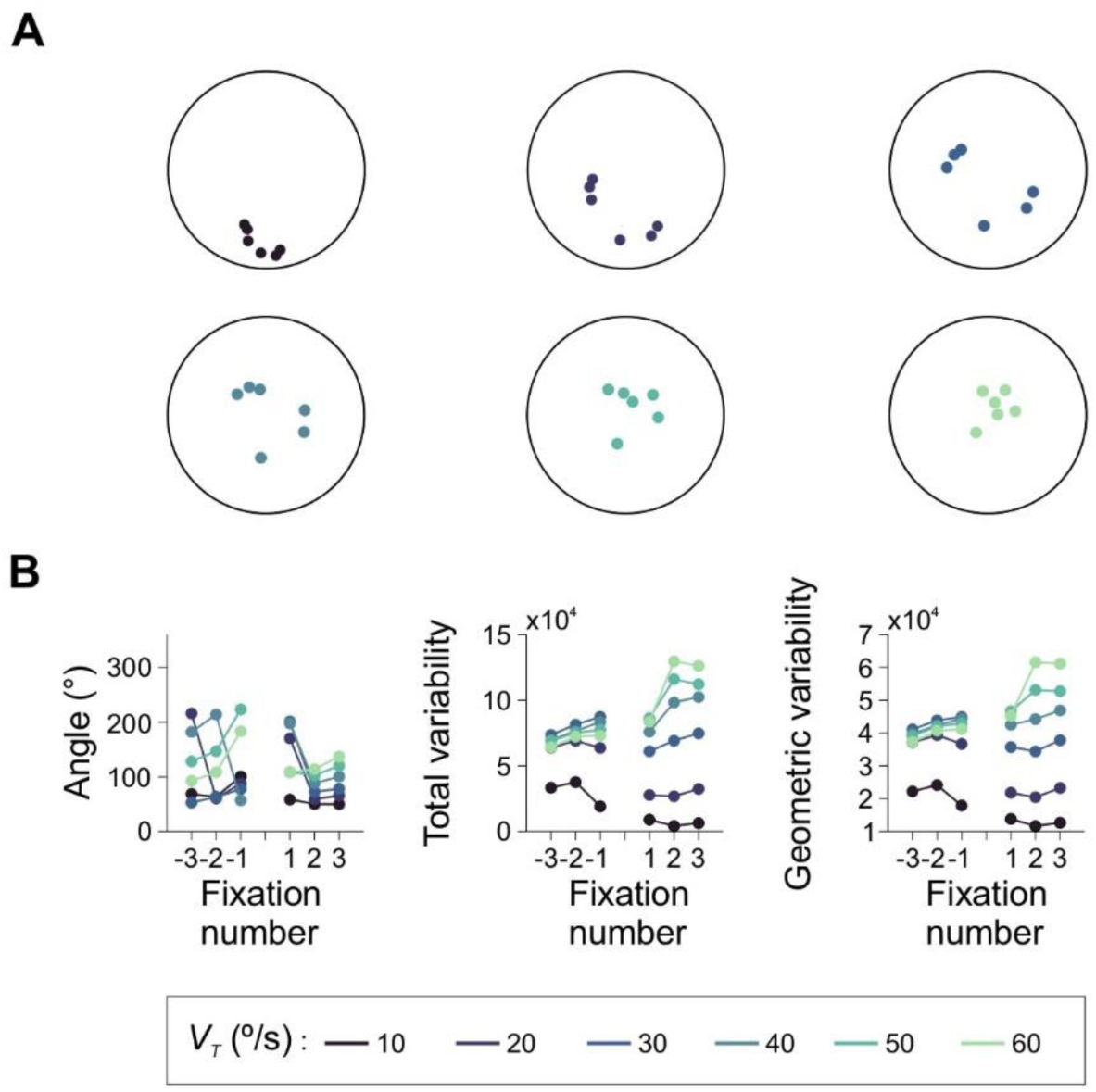
Impact of target speed on Fixation Distributions before and After Rebounds. *A*: Centers of mass for fixation distributions across three fixations before and three fixations after the rebound. Colorscale represents different target speeds, illustrating how speed influences the spatial location of fixations. Slower target speeds were associated with fixations closer to the arena walls, while higher speeds shifted fixations farther from the walls, indicating an adaptation to speed dynamics. *B*: Left panel: average orientations of fitted ellipsoids to fixation distributions as a function of fixation number, revealing how orientation shifts with target speed. Center and right panels: average total and geometric variability of fixation distributions as a function of fixation number, showing how variability is modulated by speed around rebounds.

Target speed did not affect fixation distribution angles before rebounds (Pearson correlation, *r* = 0.982, *P* = 0.12; Spearman’s rank-order correlation, *ρ* = 1, *P* = 0.33). However, after rebounds, fixation distribution angles showed improved alignment with the target’s trajectory at higher speeds (Pearson correlation, *r* = 0.984, *P* = 0.01; Spearman’s rank-order correlation, *ρ* = 1, *P* = 0.33). Two-sample Kolmogorov-Smirnov tests comparing ΔSDE values for current and future target positions across speeds yielded significant differences (*P* < 0.0001 for all cases), demonstrating participants’ refined fixation distributions and enhanced predictive strategies for all *v_T_* tested (**Figure 5B**).

### Masking the target during rebounds alters predictive fixations

To explore the impact of visual feedback on predictive fixations, we analyzed the effects of masking either the target or the user. Previous studies emphasize the importance of target visibility on anticipatory saccades (Mann et al. 2019; Márquez and Treviño 2024b). We masked either the user’s or the target’s dot by switching their contrast to 0%, making them invisible (Márquez and Treviño 2024b). This occlusion was applied at various randomly permuted inter-dot distances (IDD = 0, 3, 5, 10, corresponding to 0°, 0.8°, 1.4°, and 2.7°).

As expected, fixation distributions were disrupted when the target was masked, resulting in increased inter-dot distance (IDD) and gaze-to-target distance (GTD) values (left panels of **Figure 6A**). Target masking impaired predictive fixations, increasing GTD and reducing spatial accuracy. Heatmaps showed diminished alignment with future target positions at closer masking distances, while masking at farther distances allowed participants to rely on previously acquired trajectory information (right panels of **Figure 6A**). Statistical analyses confirmed these disruptions. A repeated-measures ANOVA revealed significant effects of masking distances on ΔSDE (*P* < 0.0001) and systematic differences in fixation distributions across fixation numbers. Kolmogorov-Smirnov tests indicated significant deviations in fixation patterns under masked versus unmasked conditions (*P* < 0.0001).

**Figure 6.**
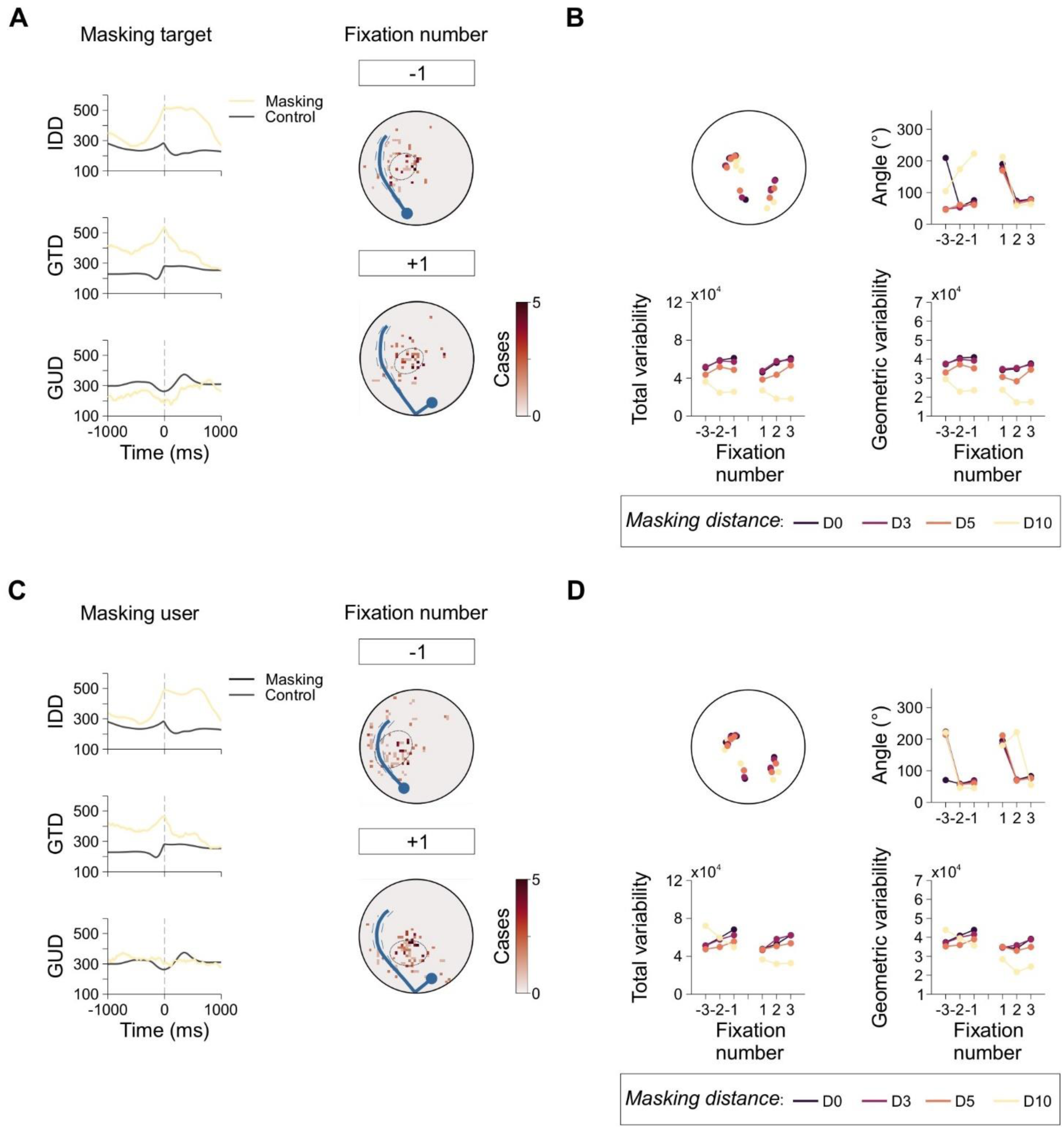
Impact of Target and User Masking on Fixation Distributions Before and After Rebounds. *A*: Event-triggered averages for inter-dot distance (IDD), gaze-to-target distance (GTD), and gaze-to-user distance (GUD) during target masking at a distance of D = 10 (see **Methods**). Black traces represent control data, while colored traces show the effect of masking. Heatmaps to the right display fixation distributions under masking, revealing substantial disruptions compared to the control distributions (from Figure 1). *B*: Insets depict the center of mass of fixation distributions, orientation angles of ellipsoids fitted to fixation distributions, and total and geometric variability as functions of fixation number. The color legend below indicates masking conditions. *C,D*: Similar analyses as in (*A*) and (*B*) but for user masking instead of target masking.

Tests involving user masking showed reduced alignment primarily at closer distances, but without significant impacts on fixation variance at greater masking distances (**Figure 6C**). A two-sample Kolmogorov-Smirnov test comparing ΔSDE between fixation positions (-3 to 3) and current and future target positions (forward frame nr 14) revealed significant differences across all masking conditions tested (*P* < 0.0001 for all cases). These results affirm the critical role of continuous visual feedback about the target in maintaining precise predictive fixations.

## DISCUSSION

This study explored predictive fixations during a visuomotor interception task, where participants used a joystick to intercept a target bouncing within a circular arena. We employed a threshold-based classification system to categorize eye movements into saccades, smooth pursuits, and fixations, ensuring reliable differentiation under dynamic task conditions. Saccades were identified by velocity and acceleration thresholds, smooth pursuits by their alignment with target speeds, and fixations as periods of gaze stability. While alternative methods, such as angular dispersion or Kalman filters, exist, our approach offered a robust classification well suited to the demands of our task.

When solving our task, participants coordinated their gaze and manual responses effectively and showed anticipatory behaviors aligned with the target’s trajectory. We analyzed gaze, joystick, and target trajectories around rebounds of the moving target using a standardized reference system. Our analyses showed that joystick positions and gaze fixations consistently aligned with the target’s future path, demonstrating participants’ ability to anticipate motion rebound trajectories.

Predictive movement timing arises from the interplay between internal forward models and external sensory feedback. Forward models compare expected and actual outcomes to detect errors rapidly, while sensory cues provide real-time feedback for corrections. This balance supports adaptive responses to dynamic environments, as observed in the rebound processing of our task and others (Badler and Heinen 2006; Kim et al. 2022; Limanowski et al. 2017).

To assess predictive behavior, we calculated squared distance errors (SDEs) between gaze positions and the target’s current and future locations. Negative ΔSDE values, indicating closer alignment with future target positions, served as markers of predictive behavior, while positive values indicated more reactive responses. Our analyses revealed negative ΔSDE values across fixation numbers, suggesting predictive strategies. These findings align with prior research, where negative ΔSDE values also reflected enhanced anticipatory responses (Brand et al. 2024; Brenner and Smeets 2011; Márquez and Treviño 2024b). Other metrics that assess predictive behavior by analyzing fixation locations in relation to target trajectories have been employed to study predictive control mechanisms (Brand et al. 2024; Müller and Sternad 2003). Evidence from areas such as object manipulation demonstrates the broader implications of predictive control. Research on object manipulation indicates that grip force profiles are shaped by anticipatory mechanisms, such as predicting an object’s weight using visual cues (Buckingham et al. 2009; Hermsdörfer et al. 2011; Mon-Williams and Murray 2000). In a similar way, tasks involving oscillatory movements demand precise adjustments to dynamic forces, illustrating the complexity involved in integrating sensory and motor predictions under varying timing conditions (Wolpert and Flanagan 2001).

Our temporal analysis of eye movement onset times relative to rebounds revealed anticipatory changes in fixations, smooth pursuits, and saccades. Peri-event time histograms (PETHs) indicated temporal shifts in gaze onset toward rebounds, supporting the role of predictive mechanisms in gaze behavior. Predictive fixations directed the gaze ahead of the target to post-rebound locations before rebounds, while saccadic activity realigned the gaze with the new trajectory afterward. This adaptive modulation demonstrates shared mechanisms between saccades and fixations, highlighting the visual system’s role in coordinating sensorimotor performance (Anderson et al. 2008; Barnes 2008; Krauzlis 2004; Krauzlis et al. 2017; Maruta et al. 2013; Orban de Xivry and Lefèvre 2007).

Our analysis of the entry and exit angles of the rebounding target revealed the impact of trajectory angles on predictive behavior. Fixations consistently aligned with the target’s future positions across all tested angles, with strong correlations between entry angles and post-rebound fixations. However, exit angles had minimal influence, suggesting that post-rebound gaze adjustments relied more on immediate trajectory information than on exit direction. These findings highlight the importance of directional cues in shaping predictive gaze strategies while also revealing specific adjustments to the unique dynamics of the task.

We also discovered that target speed influenced predictive strategies, with higher speeds resulting in fixations that were farther ahead. Heatmap analyses revealed that faster target velocities led to shifts in fixation patterns closer to predicted rebound points. Higher speeds were associated with predictive adjustments, as participants positioned their fixations closer to rebound and exit points as target speed increased. These findings demonstrate that participants adapted their visual strategies to maintain anticipatory control under heightened temporal demands.

Masking experiments clarified the role of visual feedback for predictive fixations in our task. Masking the target disrupted predictive gaze behavior, increasing gaze-to-target distance (GTD), reducing accuracy, and increasing variability. Conversely, masking the user had minimal impact, indicating that participants effectively compensated by solely relying on target trajectory cues. These results emphasize the critical role of continuous visual information about the target in maintaining predictive strategies while suggesting that self-position feedback plays a less essential role in predictive tracking (Khan et al. 2005; Márquez and Treviño 2024b).

Predictive mechanisms utilize past experiences, such as recognizing behavioral patterns in opponents, and innate physical knowledge, like understanding gravity’s effects (McIntyre et al. 2001). These mechanisms become vital when sensory input is limited, delayed, or insufficient for real-time adjustments. By enabling preemptive motor planning, predictive control ensures successful interaction despite incomplete sensory feedback. In object interaction tasks, predictive processing allows for continuous coordination with the object’s behavior. In high-pressure sports scenarios, predictive control is indispensable for timely and accurate responses under extreme temporal constraints. For example, during football penalty kicks, where the ball reaches the goal in 300–450 ms, goalkeepers rely heavily on expectations based on the kicker’s tendencies and body posture to decide their diving direction (Causer et al. 2017; Timmis et al. 2014). Conversely, goalkeepers can incorporate visual ball motion into their predictions in free kicks, where the ball takes over 1000 ms to reach the goal. Proposed strategies include predicting the ball’s position at a specific future moment or even forecasting its entire trajectory to determine an interception point. While both possibilities align with motor programming theories, conclusive evidence remains scarce, emphasizing the need for further investigation (Brenner and Smeets 2011). Expert athletes in other sports, such as volleyball players who anticipate ball direction by analyzing the setter’s hand, leg, and trunk movements, employ predictive control by effectively encoding these sport-specific signals (Piras et al. 2014).

Predictive saccades reflect long-term adaptation, with their timing influenced by prior trials, suggesting the refinement of internal models through learning. Expertise further enhances predictive control, enabling experienced individuals to adopt efficient eye movement strategies that reduce reliance on reactive adjustments. Such knowledge enables smoother and more accurate target tracking in dynamic tasks, particularly during rapid or unpredictable events. While reactive saccades are essential for immediate corrections to sudden positional changes (Chen-Harris et al. 2008), predictive saccades anticipate motion paths, facilitating anticipatory tracking (Česonis and Franklin 2020; Márquez and Treviño 2024b). As internal prediction errors diminish through practice and adaptation, reliance on predictive saccades increases, improving tracking precision and task performance (Chen-Harris et al. 2008).

Predictive control in interception tasks presumably integrates sensory feedback with forward models to optimize motor responses. Continuous visual feedback refines motor outputs by allowing participants to minimize sensory prediction errors and adapt to real-time task demands, ensuring precise alignment with target trajectories and enabling ongoing corrections to maintain accuracy (Desmurget and Grafton 2000; Limanowski et al. 2017; Miall et al. 1985; Straub and Rothkopf 2024). Forward models generate internal representations by integrating sensory and non-sensory signals, facilitating adaptable and precise motor responses even in unpredictable environments (Desmurget and Grafton 2000).

The interplay between predictive and reactive strategies has been examined in dynamic tasks. For example, in a visually guided falling-dot task, we explored how interception behaviors were influenced by manipulating external factors such as horizontal velocity, gravity, and air friction (Márquez et al. 2024). Low target variability favored predictive strategies, while high variability increased reliance on more reactive responses guided by real-time feedback. Interestingly, although manual responses showed greater variability, participants maintained stable interception performance across sessions, reflecting individual tendencies toward predictive or reactive strategies. These findings illustrate the flexibility of sensorimotor control, where anticipatory and feedback-driven mechanisms coexist to enhance performance in dynamic environments.

The brain’s motor control hierarchy relies on visual feedback to compute prediction errors and execute corrections. The posterior parietal cortex (PPC) plays a critical role in evaluating visuomotor congruence and comparing expected and actual sensory input to guide adjustments. Within the PPC, the medial region integrates visual and motor information to encode spatial trajectories. This process uses eye movement signals to perform precise visuomotor transformations, enabling the brain to predict and respond to dynamic changes in the environment (Breveglieri et al. 2012). Consistent target trajectories strengthen internal models, enhancing predictive accuracy by reducing reliance on corrective adjustments. Conversely, unpredictable trajectories disrupt this alignment, requiring greater cognitive effort for recalibration and adaptation (Bond and Taylor 2015). Similarly, the lateral occipitotemporal cortex processes early visual prediction errors, forming a feedback loop for real-time adjustments to unpredictable target movements (Limanowski et al. 2017). Computational models further elucidate the importance of feedback information in motor control. Straub and Rothkopf (2024) compared the constant bearing angle (CBA) strategy, which relies on reactive adjustments, with a Bayesian model-based approach incorporating internal predictions. While both models performed similarly under ideal conditions, the Bayesian model proved more robust under realistic conditions, such as perceptual uncertainty and sensorimotor delays, emphasizing the importance of internal models for stable motor behaviors (Straub and Rothkopf 2024).

Distinguishing predictive from prospective control illustrates the complexity of sensorimotor adaptation. Predictive control anticipates future events based on current information, while prospective control adjusts real-time actions to ongoing changes. Both strategies rely on continuous visual input to refine actions during interceptive tasks (Brenner and Smeets 2011; Márquez and Treviño 2024b). Adaptive behaviors likely integrate predictive and reactive elements to optimize performance in unpredictable environments (Katsumata and Russell 2012; Márquez et al. 2024). Task complexity and individual expertise further shape the balance between these strategies, demonstrating their adaptability (Fooken et al. 2016; Müller and Abernethy 2006; Straub and Rothkopf 2024).

Studies on reactive and predictive saccades have explored this adaptive balance. Shelhamer and Joiner (2003) showed that at low target frequencies (0.2–0.3 Hz), saccades lagged target motion, reflecting reactive tracking. However, saccades became predictive at higher frequencies (0.9 Hz and above), anticipating target motion. This abrupt transition resembles phase shifts in bistable systems, illustrating the brain’s ability to rapidly switch between neural pathways for enhanced tracking performance (Shelhamer and Joiner 2003).

### Conclusions

Our task and methods have significant potential implications. In clinical settings, systematic errors in predictive fixations could aid in diagnosing cranial nerve dysfunction, while automated visuomotor tasks could complement traditional examinations by providing quantifiable assessments. Predictive modeling tasks may also enhance rehabilitation strategies for individuals with autism or age-related cognitive decline, emphasizing anticipatory control to improve functional independence. In robotics and human-computer interaction, these findings could guide the creation of systems capable of adaptive and predictive responses in dynamic environments.

Predictive saccades rely on internal models refined through prior experiences, enabling smoother and more efficient tracking of moving targets. This anticipatory ability reduces reliance on reactive adjustments, optimizing visuomotor coordination in dynamic tasks. As participants gain expertise, prediction errors decrease, leading to greater reliance on predictive saccades and improved tracking precision. The interaction between reactive and predictive strategies reveals the brain’s ability to integrate immediate corrections with anticipatory adjustments, maintaining accuracy and flexibility in dynamic environments.

## DATA AVAILABILITY

Datasets are publicly available as tab-delimited text files at https://osf.io/826vj/?view_only=056c0e766933461691dfda407d01155a. This format was chosen for its accessibility, as it does not require specialized software licenses or familiarity with proprietary storage schemes, such as MATLAB structures. A README text file within the repository details the file naming conventions.

## DISCLOSURES

### Author note

NM (code 217590936), FH (code 218482312), and AB (code 220283653) were undergraduate students in the Bachelor’s program in Chemistry, Pharmacy, and Biology at the Centro Universitario de la Ciénega, Universidad de Guadalajara. They were all part of the ‘Departamento de Ciencias Médicas y de la Vida’, and were supervised by IM. FH was a recipient of the Early Research Promotion Program (FIT) 2025 A, in the Assistant Researcher modality. This program was sponsored by the ‘Coordinación de Investigación y Posgrado’, Centro Universitario de la Ciénega, Universidad de Guadalajara.

### Ethics approval

The study was approved by the local ethics committee of the Instituto de Neurociencias ethics committee, Universidad de Guadalajara, México, under the references ET102021-330 and ET122023-382.

### Consent to participate

All participants provided written informed consent to participate.

### Consent for publication

All participants provided written permission to publish anonymized data.

### Declaration of AI-Assisted Technologies in the Writing Process

We used Grammarly during the preparation of this manuscript to improve the text and resolve language-related issues stemming from writing in a non-native language. The final content was carefully reviewed and revised by all author(s) to ensure its accuracy, with full responsibility assumed for the published version.

## AUTHOR CONTRIBUTIONS

Conceptualization: MT; Data curation: IM, MT; Formal analysis: MT; Funding acquisition: MT; Investigation: IM, NM, FH, AB; Methodology: IM, MT; Project administration: IM; Resources: IM, MT; Software: MT; Supervision: IM, MT; Validation: IM, MT; Visualization: IM, MT; Writing—original draft: MT; Writing—review and editing: IM, MT.

## ACKNOWLEDGMENTS

Dr. Luis Lemus for valuable discussions during the development of the analysis. We acknowledge the reviewers for their constructive feedback, which significantly improved our manuscript.

## Bibliography

Anderson AJ, Yadav H, Carpenter RHS. Directional prediction by the saccadic system. Curr Biol 18: 614–618, 2008.

Badler JB, Heinen SJ. Anticipatory movement timing using prediction and external cues. J Neurosci 26: 4519– 4525, 2006.

Barnes GR. Cognitive processes involved in smooth pursuit eye movements. Brain and Cognition 68: 309–326, 2008.

Bond KM, Taylor JA. Flexible explicit but rigid implicit learning in a visuomotor adaptation task. Journal of Neurophysiology 113: 3836–3849, 2015.

Brand TK, Schütz AC, Müller H, Maurer H, Hegele M, Maurer LK. Sensorimotor prediction is used to direct gaze toward task-relevant locations in a goal-directed throwing task. J Neurophysiol 132: 485–500, 2024.

Brenner E, Smeets JBJ. Continuous visual control of interception. Hum Mov Sci 30: 475–494, 2011.

Breveglieri R, Hadjidimitrakis K, Bosco A, Sabatini SP, Galletti C, Fattori P. Eye Position Encoding in Three-Dimensional Space: Integration of Version and Vergence Signals in the Medial Posterior Parietal Cortex. J Neurosci 32: 159–169, 2012.

Buckingham G, Cant JS, Goodale MA. Living in a material world: how visual cues to material properties affect the way that we lift objects and perceive their weight. J Neurophysiol 102: 3111–3118, 2009.

Causer J, Smeeton NJ, Williams AM. Expertise differences in anticipatory judgements during a temporally and spatially occluded task. PLoS One 12: e0171330, 2017.

Česonis J, Franklin DW. Time-to-Target Simplifies Optimal Control of Visuomotor Feedback Responses. eNeuro 7: ENEURO.0514-19.2020, 2020.

Chen-Harris H, Joiner WM, Ethier V, Zee DS, Shadmehr R. Adaptive Control of Saccades via Internal Feedback. J Neurosci 28: 2804–2813, 2008.

Coen-Cagli R, Coraggio P, Napoletano P, Schwartz O, Ferraro M, Boccignone G. Visuomotor characterization of eye movements in a drawing task. Vision Res 49: 810–818, 2009.

Desmurget M, Grafton S. Forward modeling allows feedback control for fast reaching movements. Trends Cogn Sci 4: 423–431, 2000.

Faul F, Erdfelder E, Lang A-G, Buchner A. G*Power 3: a flexible statistical power analysis program for the social, behavioral, and biomedical sciences. Behav Res Methods 39: 175–191, 2007.

Fooken J, Yeo S-H, Pai DK, Spering M. Eye movement accuracy determines natural interception strategies. J Vis 16: 1, 2016.

Gould IC, Rushworth MF, Nobre AC. Indexing the graded allocation of visuospatial attention using anticipatory alpha oscillations. J Neurophysiol 105: 1318–1326, 2011.

Henderson JM. Gaze Control as Prediction. Trends Cogn Sci 21: 15–23, 2017.

Henriques DYP, Klier EM, Smith MA, Lowy D, Crawford JD. Gaze-Centered Remapping of Remembered Visual Space in an Open-Loop Pointing Task. J Neurosci 18: 1583–1594, 1998.

Hermsdörfer J, Li Y, Randerath J, Goldenberg G, Eidenmüller S. Anticipatory scaling of grip forces when lifting objects of everyday life. Exp Brain Res 212: 19–31, 2011.

Itti L, Koch C. A saliency-based search mechanism for overt and covert shifts of visual attention. Vision Res 40: 1489–1506, 2000.

Joch M, Hegele M, Maurer H, Müller H, Maurer LK. Brain negativity as an indicator of predictive error processing: the contribution of visual action effect monitoring. J Neurophysiol 118: 486–495, 2017.

Katsumata H, Russell DM. Prospective versus predictive control in timing of hitting a falling ball. Exp Brain Res 216: 499–514, 2012.

Khan AZ, Pisella L, Rossetti Y, Vighetto A, Crawford JD. Impairment of gaze-centered updating of reach targets in bilateral parietal-occipital damaged patients. Cereb Cortex 15: 1547–1560, 2005.

Kim OA, Forrence AD, McDougle SD. Motor learning without movement. Proc Natl Acad Sci U S A 119: e2204379119, 2022.

Koch I, Keller P, Prinz W. The Ideomotor approach to action control: Implications for skilled performance. International Journal of Sport and Exercise Psychology 2: 362–375, 2004.

Krauzlis RJ. Recasting the smooth pursuit eye movement system. J Neurophysiol 91: 591–603, 2004.

Krauzlis RJ, Goffart L, Hafed ZM. Neuronal control of fixation and fixational eye movements. Philos Trans R Soc Lond B Biol Sci 372: 20160205, 2017.

Land MF, McLeod P. From eye movements to actions: how batsmen hit the ball. Nat Neurosci 3: 1340–1345, 2000.

Limanowski J, Kirilina E, Blankenburg F. Neuronal correlates of continuous manual tracking under varying visual movement feedback in a virtual reality environment. Neuroimage 146: 81–89, 2017.

Liversedge SP, Gilchrist I, Everling S, editors. The Oxford Handbook of Eye Movements. Oxford University Press.

Mann DL, Nakamoto H, Logt N, Sikkink L, Brenner E. Predictive eye movements when hitting a bouncing ball. J Vis 19: 28, 2019.

Márquez I, Lemus L, Treviño M. A continuum from predictive to online feedback in visuomotor interception. Eur J Neurosci, 2024. doi:10.1111/ejn.16628.

Márquez I, Treviño M. Pupillary responses to directional uncertainty while intercepting a moving target. R Soc Open Sci 11: 240606, 2024a.

Márquez I, Treviño M. Visuomotor predictors of interception. Plos One, 2024b. doi:10.1371/journal.pone.0308642.

Maruta J, Heaton KJ, Kryskow EM, Maule AL, Ghajar J. Dynamic visuomotor synchronization: quantification of predictive timing. Behav Res Methods 45: 289–300, 2013.

McIntyre J, Zago M, Berthoz A, Lacquaniti F. Does the brain model Newton’s laws? Nat Neurosci 4: 693–694, 2001.

Mennie N, Hayhoe M, Sullivan B. Look-ahead fixations: anticipatory eye movements in natural tasks. Exp Brain Res 179: 427–442, 2007.

Miall RC, Weir DJ, Stein JF. Visuomotor tracking with delayed visual feedback. Neuroscience 16: 511–520, 1985.

Mon-Williams M, Murray AH. The size of the visual size cue used for programming manipulative forces during precision grip. Exp Brain Res 135: 405–410, 2000.

Müller H, Sternad D. A randomization method for the calculation of covariation in multiple nonlinear relations: illustrated with the example of goal-directed movements. Biol Cybern 89: 22–33, 2003.

Müller S, Abernethy B. Batting with occluded vision: an in situ examination of the information pick-up and interceptive skills of high- and low-skilled cricket batsmen. J Sci Med Sport 9: 446–458, 2006.

Orban de Xivry J-J, Lefèvre P. Saccades and pursuit: two outcomes of a single sensorimotor process. J Physiol 584: 11–23, 2007.

Pfeuffer CU, Kiesel A, Huestegge L. A look into the future: Spontaneous anticipatory saccades reflect processes of anticipatory action control. J Exp Psychol Gen 145: 1530–1547, 2016.

Piras A, Lobietti R, Squatrito S. Response time, visual search strategy, and anticipatory skills in volleyball players. J Ophthalmol 2014: 189268, 2014.

Sailer U, Flanagan JR, Johansson RS. Eye–Hand Coordination during Learning of a Novel Visuomotor Task. J Neurosci 25: 8833–8842, 2005.

Shalom DE, Dagnino B, Sigman M. Looking at Breakout: urgency and predictability direct eye events. Vision Res 51: 1262–1272, 2011.

Shelhamer M, Joiner WM. Saccades Exhibit Abrupt Transition Between Reactive and Predictive, Predictive Saccade Sequences Have Long-Term Correlations. Journal of Neurophysiology 90: 2763–2769, 2003.

Straub D, Rothkopf C. If it looks like online control, it is probably model-based control [Online]. Proceedings of the Annual Meeting of the Cognitive Science Society 46, 2024https://escholarship.org/uc/item/0gk118nw [9 Dec. 2024].

Taylor JA, Ivry RB. Flexible Cognitive Strategies during Motor Learning. PLoS Comput Biol 7: e1001096, 2011.

Timmis MA, Turner K, van Paridon KN. Visual search strategies of soccer players executing a power vs. placement penalty kick. PLoS One 9: e115179, 2014.

Treviño M, Castiello S, Arias-Carrión O, Torre-Valdovinos BD la, León RMC y. Isomorphic decisional biases across perceptual tasks. PLOS ONE 16: e0245890, 2021.

Treviño M, Medina-Coss y León R, Támez S, Beltrán-Navarro B, Verdugo J. Directional uncertainty in chase and escape dynamics. Journal of Experimental Psychology: General 153: 418–434, 2024.

Wolpert DM, Flanagan JR. Motor prediction. Curr Biol 11: R729-732, 2001.

